# Force-regulated catch bonds and fusion peptide exposure drive coronavirus entry

**DOI:** 10.64898/2026.05.21.727024

**Authors:** Haiyue Li, Zhenhai Li, Huajian Gao

**Affiliations:** Shanghai Key Laboratory of Mechanics in Energy Engineering, Shanghai Institute of Applied Mathematics and Mechanics, School of Mechanics and Engineering Science, Shanghai University, Shanghai 200072, China; Mechano-X Institute, Applied Mechanics Laboratory, Department of Engineering Mechanics, Tsinghua University, Beijing 100084, China

**Keywords:** coronavirus, catch bond, viral invasion, force regulation

## Abstract

Coronaviruses invade human cells within dynamic mechanical environments through endocytosis and membrane fusion, both mediated by the class I fusion protein spike. In SARS-CoV and SARS-CoV-2, the spike engages the human ACE2 receptor through a catch bond—an interaction whose lifetime increases under tensile force. Concurrently, mechanical pulling facilitates disruption of the S1/S2 subunits of spike, a critical step for membrane fusion. To elucidate how mechanical cues coordinate these processes, we developed a unified elastic-stochastic model that integrates theoretical analysis and computational simulations to trace viral entry. Our results identify the force-regulated catch bond between spike and ACE2 as a key determinant of successful invasion. This catch bond not only enhances receptor-mediated endocytosis but also increases the probability of S1/S2 disengagement, thereby promoting membrane fusion. Importantly, under conditions of strong catch bonding, the force-accelerated separation of S1 and S2 fine-tunes the balance between entry pathways. These findings uncover a potential mechanobiological mechanism that mediates viral cell entry by coupling receptor binding strength with spike disassembly under force. By characterizing these mechanical regulations, this work facilitates the assessment of emerging viral threats and inspires the design of drug delivery systems that leverage catch-bond kinetics for enhanced targeting.

## Introduction

Coronaviruses are a family of viruses that infect mammals and birds. To date, seven coronaviruses have been identified that infect humans, three of which—severe acute respiratory syndrome coronavirus (SARS-CoV), severe acute respiratory syndrome coronavirus 2 (SARS-CoV-2), and Middle East respiratory syndrome coronavirus (MERS-CoV)—are widely spread and can cause severe, sometimes fatal, respiratory infections^1^. SARS-CoV and MERS-CoV have led to several thousand confirmed human cases, with high mortality rates of 9.5% and 33.4%, respectively^1,2^. In contrast, SARS-CoV-2 has caused the global COVID-19 pandemic, with over 700 million infections reported to date^3^. Its high mutation rate has produced a large number of variants^2,4^, and its potential for cross-species transmission raises concerns about the emergence of novel human-infecting coronaviruses in the future—posing ongoing challenges to global public health^5^.

Coronavirus enters the respiratory system through inhaled air. The viral particles are roughly spherical and are characterized by spike proteins protruding from their surface. The spike is a class I fusion protein composed of S1 and S2 subunits. It plays a central role in facilitating host cell invasion^6,7^. As illustrated in Fig. 1, coronaviruses and other class I fusion protein viruses, such as HIV, Ebola virus, and influenza virus, share a common invasion process^8^. Take coronaviruses as an example, the invasion process involves several key steps: 1) The virus is first recognized by host cell receptors, such as ACE2, via the receptor-binding domain (RBD) located in the S1 subunit. 2) Successive formation of spike–receptor bonds leads to an expanded contact area. Following this expansion, viral entry occurs primarily through two distinct mechanisms: endocytosis and membrane fusion^7,9,10^. In the endocytosis pathway, entry proceeds through: 3) wrapping of the host cell membrane around the virus and 4) complete engulfment of the virus into the cell. In the membrane fusion pathway^10^, a separate sequence follows the expansion of contact: 3’) disengagement of the S1 and S2 subunits, 4’) conformational changes in S2 that release structural constraints on the fusion peptide^10,11^, 5’) insertion of the fusion peptide into the host membrane, 6’) formation of a fusion pore, and 7’) release of the viral genome into the host cytoplasm^10^ (Fig. 1). Despite significant progress, the detailed mechanisms of viral invasion and the factors governing pathway selection remain incompletely understood.

**Figure 1.**
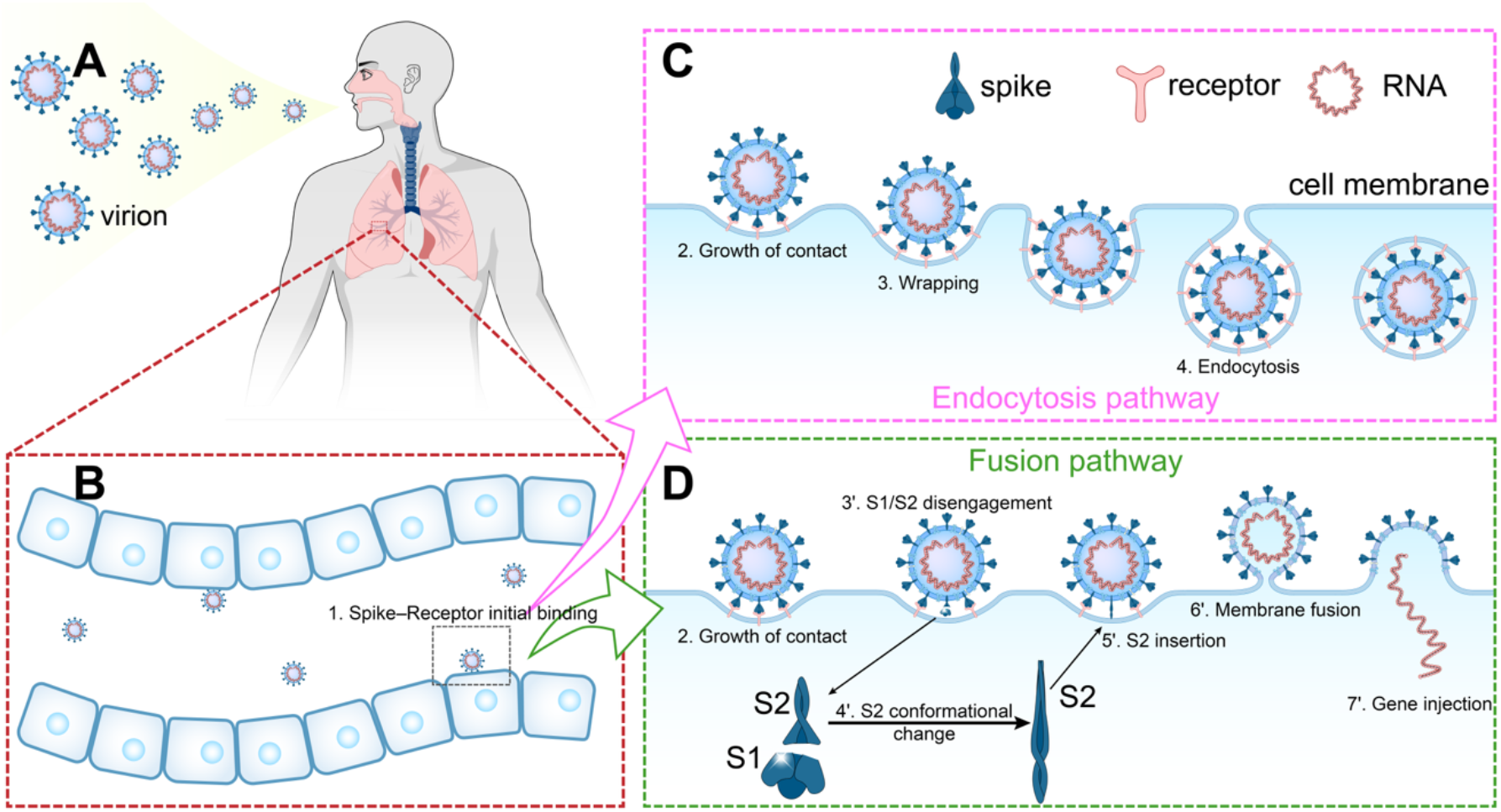
Coronavirus invasion pathways. **A**. Schematic of the mechano-environment encountered by SARS-CoV-2 during invasion of the respiratory tract. **B**. Initial binding of the virus to host cell receptors via spike–ACE2 interactions. **C**. Endocytosis pathway: progressive membrane wrapping leads to internalization of the virus into the host cell. **D**. Fusion pathway: following S1/S2 disengagement and conformational change of the S2 subunit, the viral and cellular membranes fuse, allowing viral genome entry.

Numerous studies have investigated viral endocytosis from an energetic perspective^12,13,14^. This process involves the adhesion between viral ligands and host cell receptors, where the adhesive energy generated is partially converted into elastic energy stored in the deformed cell membrane and the elongated spike–receptor bonds^12,13^. In general, endocytosis can proceed only when the binding energy released by spike–receptor interactions is sufficient to overcome the increase in elastic energy associated with membrane deformation. From a kinetic standpoint, the Arrhenius equation suggests that higher binding energy corresponds to a higher binding rate and a lower dissociation rate^15^. As such, a fast-on and slow-off spike–receptor interaction favors successful viral entry. The glycosylation of spike protein facilitates the opening of the RBD and promotes the on-rates of spike and ACE2^16,17^, which eventually might enhance the invasion. More importantly, receptor–ligand dissociation rates can be modulated by mechanical forces^18^. Rupture force analysis has been used to study the spike–ACE2 interaction, showing the loading rate could affect the disruption of spike–ACE2 bond^16,19^. Force clamping experiments have been carried out and demonstrated the human ACE2 receptor exhibits a counterintuitive catch-bond behavior when interacting with the spike proteins of SARS-CoV and SARS-CoV-2, whereby the bond lifetime increases under applied constant force^20^. During endocytosis, the spike–ACE2 bonds at the edge of the contact zone experience pulling forces, which vary based on the locations^20^. These forces, in conjunction with the catch-bond nature of the interaction, have potential to synergistically enhance viral entry. However, conventional energy-based models fall short in capturing these complex, force-dependent behaviors. In particular, they do not adequately account for the nuanced and counterintuitive effects of catch bonds on the invasion process, highlighting the need for more comprehensive mechanistic frameworks.

Moreover, the disengagement of the SARS-CoV-2 S1/S2 subunits—a prerequisite for membrane fusion—also disrupts the virus’s attachment to the host cell membrane, thereby hindering endocytosis. This S1/S2 separation is force-regulated but exhibits slip-bond characteristics, in contrast to the catch-bond behavior of the spike–ACE2 interaction; that is, force accelerates S1/S2 disengagement. As viral entry progresses, these distinct force-dependent mechanisms become tightly coupled, making the invasion process and pathway selection highly interdependent. To elucidate these dynamics, a comprehensive model that accounts for both the force-regulated spike–receptor binding and S1/S2 dissociation is essential.

To explicitly quantify the force acting on the spike–receptor bond—while accounting for both the counterintuitive catch-bond behavior of spike–ACE2 dissociation and the intuitive slip-bond nature of S1/S2 disengagement—we developed an integrated elastic–stochastic model. Since this work focuses on the force regulation, we paid minimal attention on the glycosylation and the affected on-rates of spike and ACE2. In this framework, a theoretical elastic model calculates the instantaneous force on the spike–receptor bonds during endocytosis, while a stochastic algorithm based on the kinetic Monte Carlo (KMC) method^21^ efficiently tracks the adhesion states of individual receptors and ligands on the cell membrane^22,23^. Using this model, we simulated viral invasion under various force-regulated bonding conditions and analyzed how spike–receptor reversible association and S1/S2 separation influence the entry process. Furthermore, we extended our simulations to the wild-type (WT) SARS-CoV-2 and a variant with single amino acid replacement, D614G, using experimentally determined force-regulated characteristics. The results suggest that different SARS-CoV-2 virion exhibit distinct invasion efficiencies and pathway preferences, leading to varied success rates. This study offers a new perspective on the mechanical mechanisms underlying the invasion of SARS-CoV-2 and possibly other class I fusion protein viruses, elucidating a physical mechanism describing how mechanical loading may regulate receptor-mediated adhesion and entry processes. This concept may extend to design of engineered nanoparticle delivery systems and inspire novel strategies in drug delivery.

### Concept of the Model

During the viral invasion process, spike proteins on the surface of the virus are progressively captured by host cell receptors, leading to the expansion of the virus–cell contact area and increasing curvature of the cell membrane (Fig. 2A). This membrane bending results in elevated mechanical forces exerted on the spike–receptor bonds^20,24^. These forces, in turn, modulate the stability of both the spike–receptor interaction and the S1/S2 subunit interface^20^, making the entire process highly coupled. Specifically, the dynamics of spike–receptor binding, S1/S2 disengagement, membrane deformation, and force transmission are all interdependent. To address this complex interplay, we developed an elastic–analytic model to quantitatively determine the force applied to individual bonds during membrane wrapping. In parallel, we implemented a stochastic model with Kinetic Monte Carlo (KMC) algorithm to simulate the formation and dissociation of spike–receptor and S1/S2 interactions (Fig. 2B). By integrating these two approaches, we established a unified elastic–stochastic model framework for exploring the mechanochemical feedback between binding kinetics, membrane mechanics, and force-regulated interactions. This framework provides a powerful tool to elucidate the physical principles underlying force-dependent coronavirus entry.

**Figure 2.**
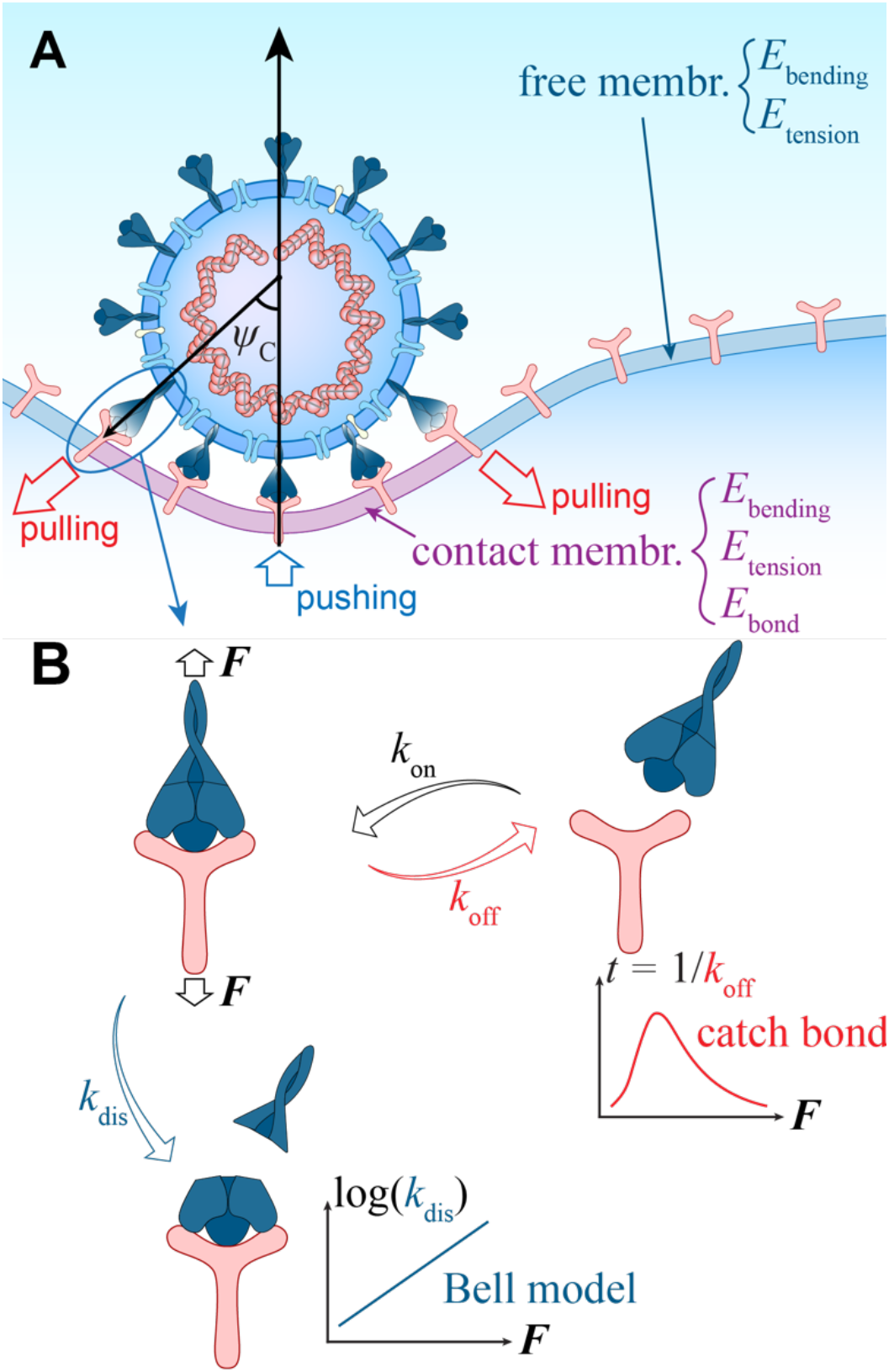
The schematic of the model concept. **A**. The analytic mechanical model. The bending and tension energy of the membrane, and the elastic energy of the spike–ACE2 bonds were considered in the model. **B**. The schematic of the kinetic model. Three groups of stochastic events—the binding of spike and ACE2, the dissociation of the spike–ACE2 bonds, and the disengagement of the S1/S2 subunits—were simulated in the KMC model. The probability of each event is governed by the transition rate: *k*_on_, *k*_off_, and *k*_dis_, respectively.

The analytical model employs two key simplifications. First, given the significant stiffness mismatch between the virus and the host cell membrane^20^, the virus is treated as undeformable. Second, the invasion is modeled as a quasi-static process consistent with prior modeling efforts^13,23^, assuming continuous mechanical equilibrium. The membrane is partitioned into contact region, where viral spikes bind to host receptors, and the free region where such interactions are absent (Fig. 2A and Fig. S1). The boundary between these states is marked by the wrapping angle, *ψ*_C_. The total energy of the system in this region comprises three components: the bending energy of the membrane, the membrane tension energy, and the elastic energy stored in the stretched spike–receptor bonds, while that in the free region includes only bending and surface tension. The equilibrium configuration is subsequently resolved by minimizing the system’s elastic energy (Supplementary Materials).

Based on the membrane configuration, we calculate the mechanical forces acting on spike–ACE2 bonds and the distances between unbound pairs. These values determine the kinetic rates: dissociation is modeled using experimental catch bond characteristics, S1/S2 disengagement via the Bell model^25^, and association through classical receptor–ligand kinetics^22,26^. These rates drive a KMC model that simulates the stochastic dynamics of binding and disengagement. At each step, an event is selected according to its transition probability; bond formation expands the contact region, while dissociation causes it to shrink. The membrane configuration is then iteratively updated based on the revised contact area (Supplementary Materials).

## Results

In order to investigate how the force regulates the virus invasion, we mainly focused on the force-regulated spike–receptor dissociation and S1/S2 disengagement with varied mechanical properties of the cell membrane and spike–ACE2 bond. To dissect the individual contributions of spike– receptor binding/dissociation and S1/S2 disengagement in mediating viral invasion, we first simulated the invasion process by considering only the stochastic binding and unbinding of spike– receptor interactions. This initial analysis allowed us to isolate the role of receptor engagement and mechanical stabilization in promoting membrane wrapping and viral uptake. We then extended the model to include the force-regulated S1/S2 disengagement, enabling us to evaluate how this additional step influences the progression of the invasion and the selection of entry pathways.

### Catch Spike–Receptor Bond Facilitates Virus Invasion

To assess the role of force-regulated spike–receptor interactions in viral invasion, we simulated the entry process under two distinct bond behaviors: slip bonds and catch-slip bonds. Both were modeled using the one-state, two-pathway framework: 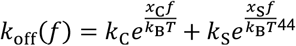, where *k*_C_, *x*_C_ and *k*_S_, *x*_S_ are the force-free dissociation rates and the corresponding transition distances of the catch- and slip-bond pathways, respectively, *k*_off_ is the force-dependent dissociation rate, *k*_R_ is the Boltzmann constant, *T* is the temperature, and *f* is the force applied to the spike–ACE2 bond (Supplementary Materials). In the simulations, the ratio *k*_C_ /*k*_S_ set to 1 for slip bonds and 1000 for catch-slip bonds, respectively. Occasional spike–receptor interactions were observed in both scenarios (Fig. S2). However, the dynamic stability and expansion of the contact area differed markedly between the two cases. In both simulations, as the virus–cell contact area grew, spike– receptor bonds at the contact edge experienced elevated pulling forces^20^ (Fig. S3). Under these conditions, slip bonds—whose lifetimes decrease with force—became unstable at the edge, limiting further growth of the contact region. In contrast, catch-slip bonds were reinforced by the pulling force, particularly at the contact edge, leading to enhanced stability. This stabilization allowed more spikes and receptors in the adjacent free membrane region to come into proximity and form new interactions, thus expanding the contact area further. This behavior established a positive feedback loop unique to catch-bond dynamics: elevated edge forces prolong bond lifetimes, which stabilize and enlarge the contact region, resulting in even higher forces at the edge. Through this mechanochemical feedback, catch-bonded spike–receptor interactions significantly promote membrane wrapping and facilitate viral entry via endocytosis (Fig. S2 and Video S1).

To further quantify the impact of force-regulated dissociation kinetics on viral invasion, we systematically varied the ratio *k*_C_/*k*_S_ from 1 to 1000 to simulate spike–receptor bonds with different catch-slip characteristics. As expected, the peak bond lifetime increased monotonically with higher *k*_C_/*k*_S_ ratios (Fig. 3A). We then incorporated these different bond types into the elastic–stochastic KMC simulations to examine their effects on virus–cell membrane interaction. The results showed a clear correlation between bond strength and membrane deformation: the maximum wrapping angle increased with the peak lifetime of the spike–receptor bond (Fig. 3B). To further evaluate the efficiency of viral entry, we defined the endocytosis efficiency as the ratio of fully wrapped viruses to the total number of simulation runs. Consistent with the enhanced mechanical stability, stronger catch bond behavior resulted in a higher endocytosis efficiency (Fig. 3C).

**Figure 3.**
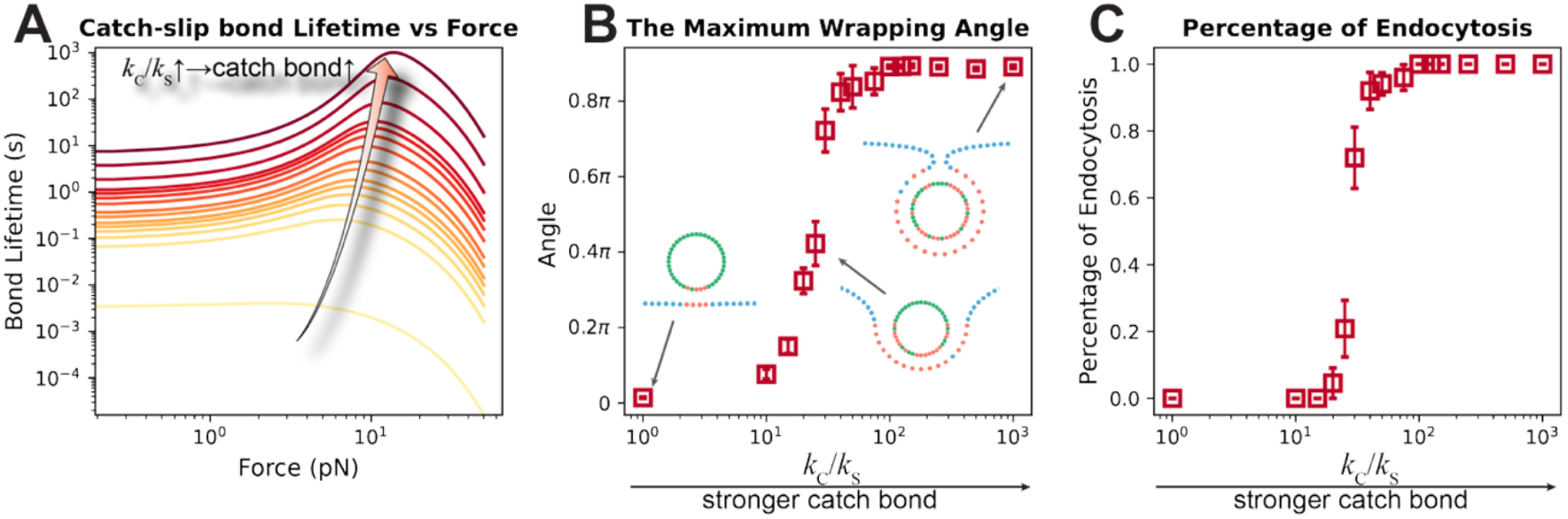
The catch spike–ACE2 bond promotes endocytosis of the virus. **A**. The catch bond characteristic is gradually enhanced by increasing *k*_C_/*k*_S_ from 1 to 1000. The inset shows that the peak lifetime of the catch bond increases monotonically with the ratio *k*_C_/*k*_S_. **B, C**. The maximum wrapping angle (**B**) and the percentage of endocytosis (**C**) during virus invasion as functions of the catch bond characteristic. In the inset of panel **B**, the green and blue dots represent the spike and ACE proteins that have not formed bonds, respectively, while the red dots represent those that have formed bonds.

### Force-Regulated S1/S2 Disengagement Regulates the Virus Invasion Pathway

To assess the role of force-regulated S1/S2 disengagement, we incorporated irreversible S1/S2 separation into the model and performed KMC simulations. A catch spike–receptor bond was used in this section to maintain growth of the contact region. The single-molecule experiments have shown the S1/S2 disengagement follows the Bell model^35,42^, 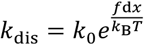, where *k*_dis_ and *k*_0_ are the force-dependent and zero-force disengagement rates, respectively, and d*x* is the characteristic transition distance along the dissociation pathway. Therefore, in the KMC simulation we modeled the distinct force regulation of S1/S2 disengagement by the Bell model with two transition distances: 0.5 nm and 4 nm for weak and strong force sensitivity, respectively. In both scenarios, the contact region initially enlarged due to the formation of spike–receptor bonds, and the catch-bond characteristics sustained longer survival times of these bonds at the edge of the contact region, which promoted further growth of the contact region. However, the stability of the spike proteins differed significantly in two conditions. In the case of weak force sensitivity of S1/S2 stability, the pulling force at the edge had limited impact on the low disengagement rate, so S1/S2 subunits rarely disengaged during invasion (Fig. S4 and Video S2). As more spike–receptor bonds formed, the cell membrane progressively wrapped around and internalized the virus. In contrast, in the case of strong force sensitivity of S1/S2 stability, the pulling force significantly increased the disengagement rate of spike proteins around the edge of the contact region. As a result, a substantial number of S1/S2 subunits disengaged. Since disengagement renders spike proteins incapable of receptor binding, the depletion of available spikes reduced receptor–ligand interactions and impaired membrane wrapping (Fig. S4 and Video S2).

To quantitively evaluate the impact of S1/S2 disengagement, we systematically modulated the force regulation by increasing the transition distance from 0.5 nm to 8 nm, while keeping the spike– receptor peak lifetime fixed at around 16 s. As expected, the sensitivity of the S1/S2 disengagement increased with a longer transition distance (Fig. 4A). We then incorporated these different force-regulated S1/S2 disengagements into the elastic–stochastic KMC simulations to examine their effects on the viral entry process. As expected, the maximum wrapping angle decreased as force-regulated disengagement became stronger (Fig. 4B). To further evaluate the likelihood of membrane fusion, we defined the disengagement percentage as the ratio of S1/S2-disengaged spikes to the total number of bound spike–receptor pairs. In contrast to the maximum wrapping angle, the disengagement percentage increased with higher force sensitivity (Fig. 4C).

**Figure 4.**
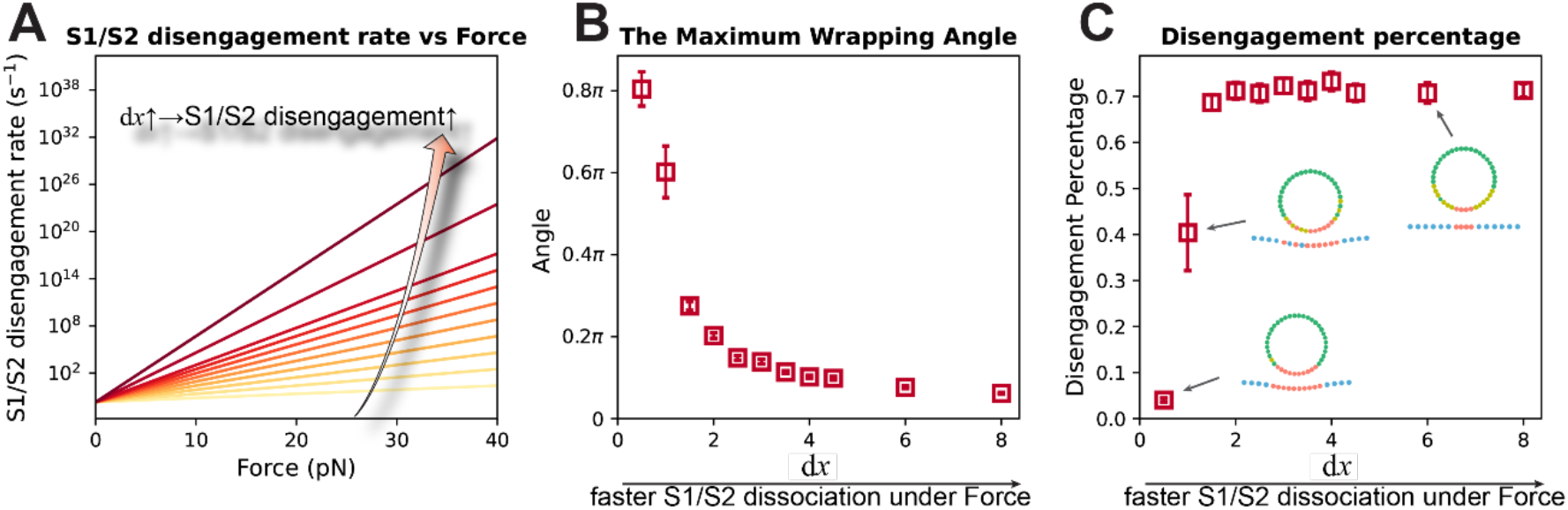
Pronounced force acceleration facilitates S1/S2 disengagement. **A**. The force acceleration of S1/S2 disengagement is increased by elevating the transition distance d*x* in the Bell model. **B, C**. The maximum wrapping angle (**B**) and the disengagement percentage (**C**) during virus invasion as functions of force acceleration on S1/S2 disengagement, evaluated at different membrane bending stiffness. Panels **B** and **C** share the same symbol definitions as in Fig. 3. In the inset of panel **C**, the green, blue, and red dots share the same definitions as in Fig. 3, while the yellow dots represent the spike protein, in which the S1/S2 has successfully disengaged..

### The Cooperation of Force-Regulated Spike–Receptor Dissociation and Force-Accelerated S1/S2 Disengagement

In the absence of S1/S2 disengagement, the catch-bond behavior of the spike–receptor interaction facilitates membrane wrapping by stabilizing the contact region under force, thereby enhancing the likelihood of endocytosis (Fig. 3B and 3C). In contrast, when S1/S2 disengagement is strongly accelerated by force, the pulling force applied to the spike promotes separation of the S1 subunit from S2 (Fig. 4), which may initiate the membrane fusion pathway.

To unravel the complicated interplay and gain a comprehensive understanding of force-regulated viral invasion, we systematically explored the key characteristics of virus wrapping and S1/S2 disengagement across a broad parameter space. Specifically, we varied the catch-slip behavior of spike–receptor bonds, ranging from a slip bond with a ∼0.004-s lifetime at zero force to a strong catch bond with a ∼1000-s peak lifetime at ∼10 pN. In parallel, we modulated the force sensitivity of S1/S2 disengagement by adjusting the transition distance from 0.5 nm to 8.0 nm. To visualize the combined effects of these parameters, we generated heatmaps of two key metrics: the disengagement percentage and the normalized maximum wrapping angle.

Viruses with weak catch spike–receptor bonds exhibit both a reduced maximum wrapping angle and a lower S1/S2 disengagement percentage (Fig. 5A and 5B). As the catch-bond behavior strengthens, the maximum wrapping angle increases, reflecting enhanced membrane wrapping due to more stable spike–receptor interactions. However, this benefit is rapidly offset when the force-acceleration of S1/S2 disengagement increases (Fig. 5A), as premature S1/S2 separation limits the number of functional spikes available for continued receptor binding. Consistent with expectations, the disengagement percentage rises significantly as the force sensitivity of S1/S2 dissociation increases, driven by larger transition distances (Fig. 5B).

**Figure 5.**
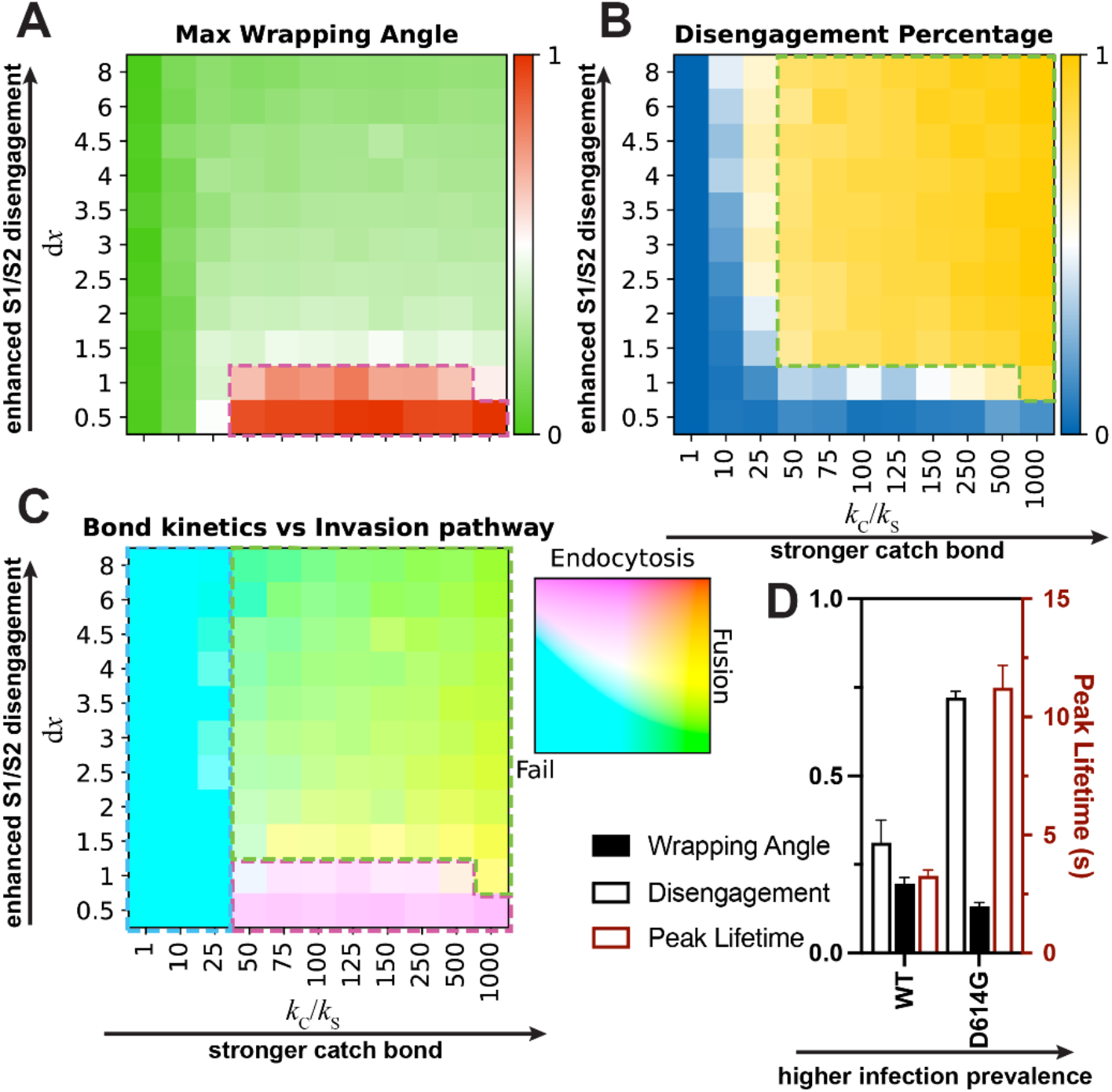
A phase diagram of virus entry features. **A, B**. Heatmaps of the normalized maximum wrapping angle (**A**) and the disengagement percentage (**B**) across the *k*_C_/*k*_S_ and d*x* parameter space, representing the extent of the spike–ACE2 catch bond and the force acceleration on S1/S2 disengagement, respectively. **C**. Phase diagram of virus invasion pathways. The disengagement percentage and the normalized maximum wrapping angle are mapped to two independent color channels of the LAB color space: yellow–blue and red–green, respectively. The blue, magenta, and green dashed boxes highlight regions where the virus: (1) fails to invade (both normalized maximum wrapping angle and disengagement percentage < 0.5), (2) favors the endocytosis pathway (wrapping angle > disengagement percentage), and (3) favors the membrane fusion pathway (disengagement percentage > wrapping angle). **D**. Peak lifetimes measured from single-molecule experiments, and model-calculated normalized maximum wrapping angles and disengagement percentages for WT and D614G variants.

## Discussion

A higher maximum wrapping angle reflects a greater likelihood of successful endocytosis. According to our simulations, increased S1/S2 disengagement impedes membrane wrapping but facilitates membrane fusion by increasing the probability of fusion peptide insertion into the host membrane^10^. By overlaying the color channels from Fig. 5A and 5B, the interplay between these two entry pathways—cooperative and competitive—can be more clearly visualized (Fig. 5C). When the spike–receptor bond behaves as a slip bond with a short force-free lifetime, viral entry fails, as neither endocytosis nor membrane fusion can be completed (blue region in Fig. 5C; failed pathway illustrated at the bottom of Fig. 6). In contrast, when the spike–receptor bond exhibits strong catch-bond behavior, the interaction is significantly prolonged under pulling forces, particularly at the contact edge. This force-enhanced stabilization enables sustained receptor engagement, allowing the virus to remain anchored and promoting progressive membrane wrapping. If S1/S2 disengagement is minimally sensitive to force, few or no S2 subunits become exposed, and fusion is unlikely to occur. In this case, endocytosis dominates, as the cell continues to internalize the virus via wrapping (violet region in Fig. 5C; endocytosis pathway in the center of Fig. 6). Conversely, when S1/S2 disengagement is strongly accelerated by force, more S2 subunits exposes. The surrounding catch spike–ACE2 bonds hold the virus and host membrane close, providing sufficient time and proximity for S2 to complete its conformational change and insert into the membrane, ultimately triggering fusion (green region in Fig. 5C; fusion pathway subset in Fig. 6). Taken together, catch-bond characteristics consistently enhance viral invasion, regardless of the entry strategy. Meanwhile, the force-regulated S1/S2 disengagement finely modulates the balance between endocytosis and membrane fusion, with the catch-bond mechanism playing a critical supporting role in both pathways. Given the shared invasion pathways among class I fusion protein viruses, force regulation in receptor–glycoprotein recognition and glycoprotein subunit disengagement may similarly influence the invasion processes of other viruses.

**Figure 6.**
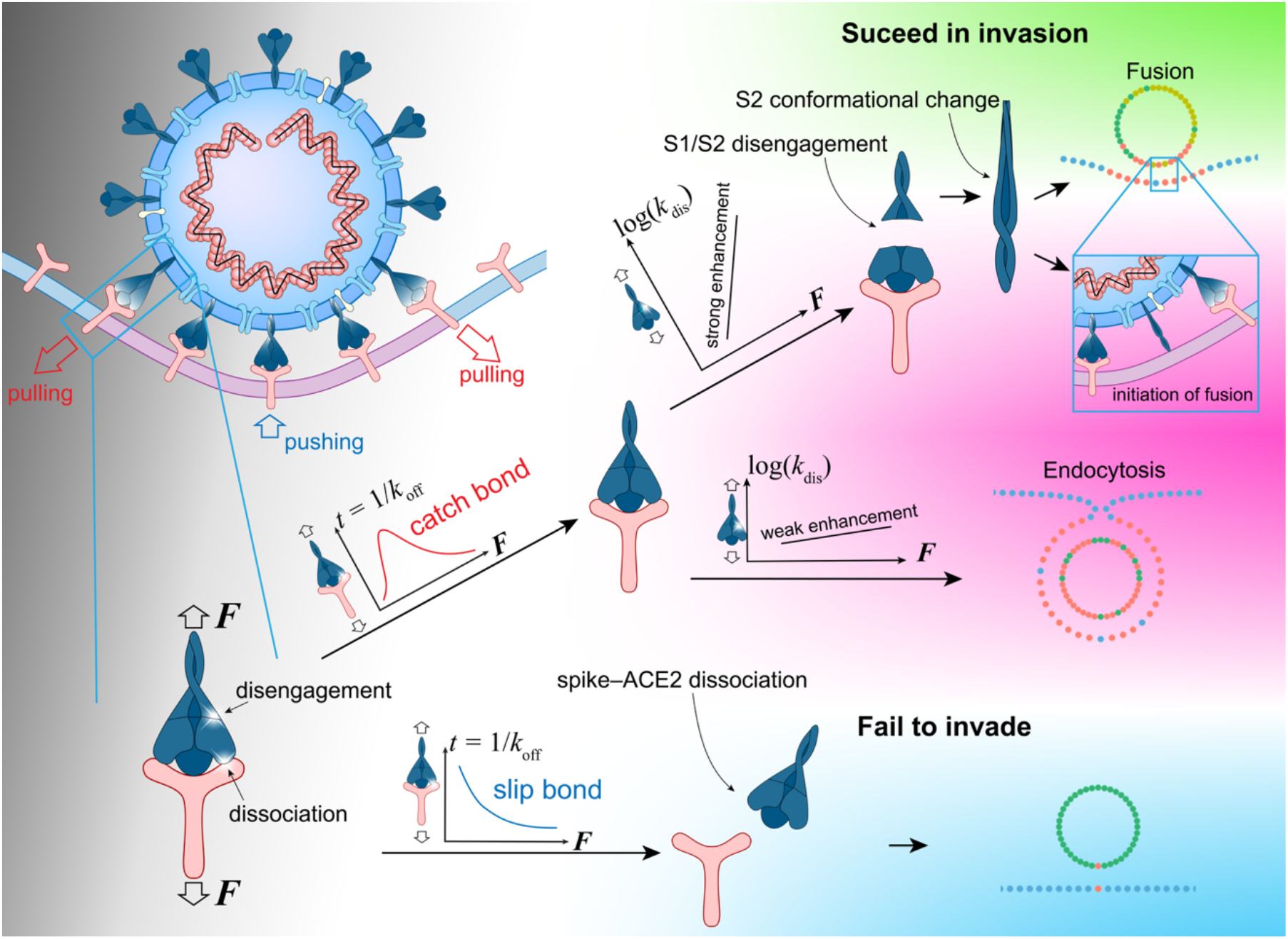
Invasion pathway selection through the cooperation between spike–ACE2 bond formation and S1/S2 disengagement. A schematic illustration of how the coordinated action of spike–ACE2 binding and force-regulated S1/S2 disengagement determines the selection between endocytosis and membrane fusion pathways during viral entry.

The widely circulating SARS-CoV-2 virus continues to evolve, giving rise to numerous variants with distinct spike protein properties. For example, the amino acid change D614G, which became dominant after April 2020^27^, exists in most major variants, including Alpha, Beta, and Delta^28^. The emergence of these variants has raised concerns regarding their infectivity and the potential role of force-regulated mechanisms in viral entry. Leveraging our model, we investigated how the WT and D614G variants interact with host cells and how force-dependent processes may vary among them. Notably, recent work by Hu et al. systematically examined the spike–ACE2 dissociation and S1/S2 disengagement behaviors of these variants using single-molecule techniques^20^. Their findings revealed that they exhibit distinct force-regulated spike–ACE2 dissociation and S1/S2 disengagement: the WT shows relatively weak catch-bond behavior, while D614G displays a stronger catch bond (Fig. 5D and Fig. S5). Interestingly, the strength of this catch bond correlates with the global prevalence of its associated WT and D614G variants (Fig. S5) ^4^. The D614G variants with higher infection rates tend to exhibit stronger catch-bond characteristics in spike–ACE2 dissociation, thereby promoting more efficient viral entry. This enhanced mechanical interaction likely contributes to the elevated infectivity of the D614G variants and its ability to outcompete the WT strain.

The D614G variant reduces the force-free S1/S2 disengagement rate^20,29^, while enhancing the force-accelerated S1/S2 disengagement behavior compared to the WT SARS-CoV-2 spike^20^. To evaluate how these force-regulated characteristics influence viral invasion and pathway selection, we incorporated the experimentally measured dissociation parameters into our model (Fig. S5 and Tables S2 and S3). Simulation results showed that, compared to WT SARS-CoV-2, the D614G variant exhibited a modest reduction in normalized maximum wrapping angle, but a substantial increase in the S1/S2 disengagement percentage (Fig. 5D). This outcome is attributed to the higher force sensitivity of S1/S2 disengagement in the variant (Fig. S5). Consistent with experimental reports, the D614G variant has been found to favor the membrane fusion pathway over endocytosis^31^, aligning with the pronounced force-accelerated S1/S2 disengagement observed in our model. These findings suggest that the enhanced force sensitivity of S1/S2 separation may contribute to a shift in the viral entry strategy, favoring fusion over endocytosis in variants with elevated infectivity.

While informative, the current model focuses primarily on molecular and mechanical interactions and does not fully capture the complexity of the cellular environment and all aspects of the viral entry process. Under physiological conditions, multiple additional steps are involved in both the endocytosis and membrane fusion pathways. First of all, the spike and ACE2 can be glycosylated at multiple position, which could affect the effective on-rate and the mechanical property of the spike–ACE2 bond. In the case of endocytosis, membrane wrapping is accompanied by several cellular activities, including the recruitment of clathrin and cargo proteins to the curved membrane to form a coated pit, the polymerization of actin filaments to provide structural support and force generation, and eventually, the constriction and scission of the membrane neck to complete vesicle internalization^32^. For the fusion pathway, following S1/S2 disengagement, the S2 subunit undergoes a substantial conformational change, which exposes the S2’ cleavage site. This site must be cleaved by the host protease TMPRSS2 to enable insertion of the fusion peptide into the host membrane and subsequent fusion of the viral and cellular membranes^10^. In the present model, we focus on elastic energy-driven membrane deformation and force-regulated protein interactions, without explicitly modeling these downstream cellular and proteolytic events, lipid rafts, endocytic machinery, and cytoskeletal factors. Therefore, the increased disengagement percentage indicates higher chances for fusion, but it’s not directly equal to the fusion percentage. To achieve a more comprehensive understanding of the viral entry dynamics and pathway selection mechanisms, future studies should integrate additional biological processes—such as glycosylation of spike and ACE2, cytoskeletal remodeling, protease activity, endocytic machinery, and the aforementioned downstream conformational changes of S2 subunit—and validate model predictions through single-molecule and live-cell imaging approaches. Expanding and refining the model in this direction will help elucidate how mechanical forces regulate viral invasion in vivo across different variants, ultimately guiding the development of more effective interventions against SARS-CoV-2 and related class I fusion protein viruses.

## Conclusion

Our simulations suggest that SARS-CoV-2 invasion is governed by the coordinated interplay between spike–ACE2 catch bonds and force-accelerated S1/S2 disengagement, which together shape both the entry efficiency and the preferred invasion pathway. By bridging molecular-scale mechanisms with cell-level outcomes, this work provides mechanistic insights that may inform the design of antiviral strategies targeting the mechanical steps of viral entry.

## Acknowledgments

This work is supported by grants from the National Natural Science Foundation of China (12272216 and 12172204) and the Natural Science Foundation of Shanghai (22ZR1423500). H.G. acknowledges support from a Tsinghua University start-up grant for the Mechano-X Institute.

## Notes

### Competing Interest Statement

The authors have declared no competing interest.

