## Supplementary Methods, Supplementary Fig. 1-5, and Supplementary Table 1, 2 for "Force-regulated catch bonds and fusion peptide exposure drive coronavirus entry"

### Supplementary Material for
#### Force-regulated catch bonds and fusion peptide exposure drive coronavirus entry

Haiyue Li[1], Zhenhai Li[1*], Huajian Gao[2*]

Huajian Gao, Zhenhai Li
.

**This PDF file includes:**

Supporting text
Figures S1 to S5
Tables S1 to S3
Legends for Movies S1 to S2
SI References

**Other supporting materials for this manuscript include the following:**

Movies S1 to S2

#### Supporting Information Text

#### Methods

##### 1. Force on the spike–receptor bonds

In the analytic model, we proposed two key assumptions: the virus remains undeformed during the invasion process, owing to the large difference in elastic modulus between the virus and the cell membrane [1]; the entire system is assumed to be quasi-static [2-4]. As the cell is orders of magnitude larger than the virus, we model the virus and cell membrane as a sphere sitting on an infinite plate. The virus and the cell membrane together form an axisymmetric system [2-4]. We build a Cartesian coordinate system $\overrightarrow{e_r}$-$\overrightarrow{e_h}$, and put the origin at the center of the virus (Fig. 2A). In our model, we partition the cell membrane into two regions: the contact region where the membrane makes close contact with the virus through spike–receptor bonds (Fig. 2A), and the free region where the receptors do not form interactions with the spikes on the virus yet. The boundary separating two regions defines the wrapping angle, $\psi_C$.

In the contact area, the deformed cell membrane can be approximated as a portion of a spherical dome with the center at $O'$ and a radius of $R$. To characterize the shape, we define relevant geometric parameters as shown in the schematic (Fig. S1). Specifically, we simplify the spike–receptor bond as a linear spring with a stiffness of $k$ and a resting length of $l_b$ [1,5]. As we consider the process to be quasi-static, the system is under equilibrium at any moment. Thus, the radius, $R$, of the contact area can be determined through energy minimization. In the equilibrated virus–cell system, the total energy of the contact area is composed of the bending energy ($E_{\text{bending}}$) and the surface tension energy ($E_{\text{tension}}$) of the deformed membrane, the bond energy ($E_{\text{bond}}$) of the stretched spike–receptor bonds.

$$E_{\text{contact}} = E_{\text{bending}} + E_{\text{bond}} + E_{\text{tension}} \tag{1}$$

The bending energy can be written as $E_{\text{bending}} = 4\pi\kappa(1-\cos\psi_C)$, where $\kappa$ is the bending stiffness. The surface tension energy can be written as $E_{\text{tension}} = \sigma\pi R^2(1-\cos\psi_C)^2$, where $\sigma$ is the surface tension, $R$ is the radius of the sphere. The total bond energy is calculated by summing the bond energies over all the spike–receptor bonds: $E_{\text{bond}} = \sum_{i=1}^{n}\frac{1}{2}k(l_i-l_b)^2$, in which $l_i$ and $l_b$ are the current and resting lengths of the $i$th spike–receptor bond, respectively. By minimizing the total contact energy $E_{\text{contact}}$, the configuration of the cell membrane within the contact area, including the radius of the cell membrane ($R$) and the gap between the apex of the virus to the cell membrane ($I_{\text{apex}}$), can be obtained. Eventually, the equilibrium length of each spike–receptor bond in the contact area can be determined by Eq. 2, while the force in the spike–receptor bond is written as Eq. 3.

$$l_i = R - a\frac{\sin\alpha_i}{\sin\alpha_i'} \tag{2}$$

$$f_i = k(l_i - l_b) \tag{3}$$

##### 2. Determination of the shape of free membrane

2.1 Energy functional and shape equations

During virus invasion, the shape of free membrane can be determined by adding the thermodynamic fluctuations on top of the equilibrium configuration obtained by minimizing the elastic energy of the system. The Fig. S1 illustrates a virus with a radius of $a$ invading a cell in the $\overrightarrow{\boldsymbol{e_r}}$-$\overrightarrow{\boldsymbol{e_h}}$ coordinate system. The current invasion angle is denoted as $\psi_C$, the local normal vector of the cell membrane is denoted as $\vec{\boldsymbol{n}}$, and the angle between the cell membrane and the $\overrightarrow{\boldsymbol{e_r}}$ direction is denoted as $\varphi$. The geometric constraints on the cell membrane shape can be written as $\dot{r} = \cos\varphi$ and $\dot{h} = \sin\varphi$ [2,4], where dot denotes the derivative with respect to arclength $s$.

The shape of free membrane in its minimum energy state is determined using the least action principle. The total energy of the free membrane is composed of the bending energy and the surface tension energy, described by the Canham-Helfrich energy [2,4]

$$E_{\text{free}} = \iint [\frac{\kappa}{2}(2H-H_0)^2 + \bar{\kappa}K + \sigma]\mathrm{d}A \tag{4}$$

where $\bar{\kappa}$ is Gaussian rigidity of the membrane, $H$ is the mean curvature, written as $\frac{1}{2}(c_1+c_2)$, and $K$ is the Gaussian curvature of the surface, written as $c_1c_2$. The represent the principal curvature of the surface $c_1$ and $c_2$ can be obtained by $c_1 = \dot{\varphi}$ and $c_2 = \sin\varphi/\mathrm{r}$, respectively. $H_0$ is the intrinsic mean curvature of the membrane, which is equal to 0 in present study. Since no topological changes is considered in our model, the

Gaussian rigidity term remains constant and is not incorporated into the model. After simplification, the total energy can be expressed as [2,6]:

$$E_{\text{free}} = \pi\kappa \int [(\dot{\varphi} + \frac{\sin\varphi}{r})^2 + \frac{2\sigma}{\kappa}(1-\cos\varphi)] r \mathrm{d}s \tag{5}$$

To further simplify the expression, we introduce two non-dimensional parameters:

$$\tilde{E} = \frac{E_{\text{free}}}{\pi\kappa}$$
$$\tilde{\sigma} = \frac{\sigma a^2}{\kappa} \tag{6}$$

In order to satisfy the geometric constraints of the free membrane, we incorporate these constraints into the total energy as Lagrange multiplier terms. Thus, the total energy can be written as [2]:

$$\tilde{E} = \int L \mathrm{d}s$$
$$L = r\left[\left(\dot{\varphi} + \frac{\sin\varphi}{r}\right)^2 + \frac{2\tilde{\sigma}}{a^2}(1-\cos\varphi)\right] + \lambda_{\text{r}}(\dot{r} - \cos\varphi) + \lambda_{\text{h}}(\dot{h} - \sin\varphi) \tag{7}$$

where $\lambda_{\text{r}}(s)$ and $\lambda_{\text{h}}(s)$ are the Lagrangian multipliers.

Apparently, the total energy of the membrane is a surface integral of local bending and tension contributions. Therefore, it is dependent on the membrane's shape. Following the least action principle, the Lagrangian function *L* should satisfy the Euler-Lagrange equation. The Lagrangian function *L* can be considered as a function of generalized coordinates ($\varphi$,$r$,$h$). The generalized momenta can be derived as [2]:

$$P_\varphi = \frac{\partial L}{\partial \dot{\varphi}} = 2r(\dot{\varphi} + \frac{\sin\varphi}{r})$$
$$P_r = \frac{\partial L}{\partial \dot{r}} = \lambda_r \tag{8}$$
$$P_h = \frac{\partial L}{\partial \dot{h}} = \lambda_h$$

These generalized momenta represent the conjugate momenta corresponding to the generalized coordinates.

As the Lagrangian *L* is independent of the arclength, *s*, the corresponding Hamiltonian is conserved. Moreover, numerically one could integrate systems of the first-order differential equations. Therefore, we switch to a Hamiltonian description. The Hamiltonian description is obtained through a Legendre transformation:

$$H = \dot{\varphi}P_\varphi + \dot{r}P_r + \dot{h}P_h - L$$
$$H = \frac{{P_\varphi}^2}{4r} - \frac{P_\varphi \sin\varphi}{r} - \frac{2\tilde{\sigma}r}{a^2}(1-\cos\varphi) + P_r\cos\varphi + P_h\sin\varphi \tag{9}$$

Then, the shape of free membrane in its minimum energy state can be obtained by solving the Hamiltonian equations with appropriate boundary conditions [2]:

$$\dot{\varphi} = \frac{\partial H}{\partial P_\varphi} = \frac{P_\varphi}{2r} - \frac{\sin\varphi}{r}$$
$$\dot{r} = \frac{\partial H}{\partial P_r} = \cos\varphi$$
$$\dot{h} = \frac{\partial H}{\partial P_h} = \sin\varphi$$
$$\dot{P_h} = -\frac{\partial H}{\partial h} = 0 \tag{10}$$
$$\dot{P_r} = -\frac{\partial H}{\partial r} = \frac{P_\varphi}{r}\left(\frac{P_\varphi}{4r} - \frac{\sin\varphi}{r}\right) + \frac{2\tilde{\sigma}}{a^2(1-\cos\varphi)}$$
$$\dot{P_\varphi} = -\frac{\partial H}{\partial \varphi} = \left(\frac{P_\varphi}{r} - P_h\right)\cos\varphi + \left(\frac{2\tilde{\sigma}r}{a^2} + P_r\right)\sin\varphi$$

2.2 Boundary conditions

Boundary condition at *s=0* is required to solve $\varphi$, $r$, $h$, $P_\varphi$, $P_r$, and $P_h$.The boundary conditions must be chosen in a way that guarantees a smooth contact at the intersection point between the free membrane and the contact area ($s$=0), while also ensuring that the membrane becomes asymptotically flat at large radial distances. These boundary conditions are given as follows [2]:

$$\begin{gathered} r(0) = R\sin\psi_C \\ \varphi(0) = \psi_C \\ h(0) = -R\cos\psi_C + R - a - I_{apex} \end{gathered} \tag{11}$$

To maintain the asymptotic flatness, we require that the angle $\varphi$ and its derivatives vanish as $s$ approaches infinity [2]:

$$\begin{gathered} \lim_{s\to\infty}\varphi(s) = 0 \\ \lim_{s\to\infty}\dot{\varphi}(s) = 0 \end{gathered} \tag{12}$$

It has been shown that the $\varphi(s)$ vanishes rapidly and any contributions beyond a large distance $S$ in arclength can be considered insignificant. Therefore, a practical way is to choose an upper arclength limit $S$ and impose the infinity condition there. In our simulation, $S$ is set to be *20a*. Moreover, additional boundary conditions at $s=S$ for $P_r$ and $P_h$ are obtained during the functional minimization [2]:

$$\begin{gathered} P_r(S) = \frac{\partial L}{\partial \dot{r}}|_{s=S} = 0 \\ P_h(S) = \frac{\partial L}{\partial \dot{h}}|_{s=S} = 0 \end{gathered} \tag{13}$$

Thus, $P_h$ must be 0 to satisfy both Eq. 10 and 13. Furthermore, the Hamiltonian $H(s)$ approaches $P_r$ as $s$ approaches infinity when assuming a flat membrane at infinity from Eq. 9. Since $H(s)$ is a conserved quantity, $H(s)$ must be 0 based on Eq. 13. Combining Eqs. 8 and 9, we derived the boundary condition for $P_r(0)$ as follows [2]:

$$RP_r(0) = \sqrt{\frac{z(2-z)}{1-z}}\left[1 + \frac{2\tilde{\sigma}R^2}{a^2}z - (R\dot{\varphi}(0))^2\right] \tag{14}$$

where $z = 1 - \cos\psi_C$. Both $P_\varphi(0)$ and $P_r(0)$ are functions of $\dot{\varphi}(0)$ according to Eqs. 8 and 14. Therefore, if $\dot{\varphi}(0)$ is given, we can easily determine $P_\varphi(0)$ and $P_r(0)$. However, it is not possible to determine the specific value of $\dot{\varphi}(0)$ at this point. To solve this problem, the shooting method has been employed. In the shooting method, a series of attempts of $\dot{\varphi}(0)$ are made, and the shape functions of the membrane are obtained by solving the Hamiltonian equations [2,4]. The set of results that best fits the boundary conditions at $S$ (Eq. 12) is then selected as the membrane shape function.

**3. Transition Rates of Bond Kinetics**

Spike–receptor binding is mediated by reversible non-covalent interactions, allowing the spike and receptor in the contact region to undergo cycles of binding, dissociation, and potential rebinding. To capture the stochastic nature of these events during viral invasion, we employed a kinetic Monte Carlo (KMC) approach, in which the formation and rupture of spike–receptor bonds, as well as the disengagement of the S1/S2 subunits, are treated as independent biochemical reaction events[5,7,8]. Each of these reactions occurs with a probability determined by its respective kinetic rate (Fig. 2B). Accurately modeling this process requires explicit definitions of the kinetic rates for all included events.

Based on magnetic tweezer experiments conducted on single spike proteins, the disengagement of the S1/S2 subunits is found to be monotonically accelerated by applied pulling forces. This behavior is characteristic of a slip bond, and the force dependence of the dissociation rate can be well described by the Bell model[1,9], $k_{\text{dis}} = k_0 e^{\frac{f\text{d}x}{k_\text{B}T}}$, where $k_{\text{dis}}$ and $k_0$ are the force-dependent and zero-force disengagement rates, respectively; $k_\text{B}$ is the Boltzmann constant, $T$ is the temperature, $f$ is the force applied to the spike protein, and d$x$ is the characteristic transition distance along the dissociation pathway. Notably, experimental observations indicate that the force-free disengagement rate is extremely low, suggesting that S1/S2 subunits rarely separate in the absence of external force[1]. In our model, we therefore assume that S1/S2 disengagement occurs only when the spike protein is simultaneously bound to the receptor and subjected to tension. Once disengagement takes place, the S2 subunit undergoes a conformational transition that may lead to its insertion into the host cell membrane, initiating membrane fusion (Fig. 1). Importantly, the dissociated S1 and S2 subunits are considered irreversibly separated and cannot rebind to reconstitute a functional spike protein. This assumption reflects the unidirectional nature of the fusion pathway and ensures that spike-mediated membrane fusion proceeds only after successful force-regulated S1/S2 dissociation.

In contrast to the S1/S2 interaction, the SARS-CoV-2 spike–ACE2 bond exhibits a strong catch-slip bond behavior, where the bond lifetime initially increases with applied force, reaches a peak, and then decreases[1,10].

To model this non-monotonic force dependence, we adopted a one-state, two-pathway kinetic framework[11,12]. In this model, bond dissociation can occur via two parallel pathways: one aligned with the pulling direction (representing the slip-bond phase), and the other opposing the pulling direction (representing the catch-bond phase). Both dissociation pathways follow the Bell model, and the overall force-dependent dissociation rate—defined as the reciprocal of the bond's mean lifetime—is given by: $k_{\mathrm{off}}(f) = k_{\mathrm{C}} e^{\frac{x_{\mathrm{C}} f}{k_{\mathrm{B}} T}} + k_{\mathrm{S}} e^{\frac{x_{\mathrm{S}} f}{k_{\mathrm{B}} T}}$[12], where $k_{\mathrm{C}}$, $x_{\mathrm{C}}$ and $k_{\mathrm{S}}$, $x_{\mathrm{S}}$ are the force-free dissociation rates and the corresponding transition distances of the catch- and slip-bond pathways, respectively. Because the catch pathway resists the pulling force, the transition distance $x_{\mathrm{C}}$ is negative, while $x_{\mathrm{S}}$ is positive. By increasing the ratio $k_{\mathrm{C}}/k_{\mathrm{S}}$ while keeping $x_{\mathrm{C}}$ and $x_{\mathrm{S}}$ constant, the strength of the catch bond effect can be modulated (Fig. 3A), allowing us to explore how catch-slip dynamics influence viral adhesion and entry efficiency.

The binding events between spike proteins and host cell receptors are governed by the association rate, $k_{\mathrm{on}}$, which depends on both the intrinsic molecular association rate and the spatial separation between the binding partners. This relationship can be described using a classical model of receptor–ligand binding kinetics[5,7,13]. According to this model, bond formation is distance-dependent and occurs only when the separation δ between the anchor points of the ligand (spike) and receptor (ACE2) falls within a predefined maximum reactive distance, denoted $l_{\mathrm{bind}}$. The binding rate, $k_{\mathrm{on}}$, depends on the virus-cell surface separation as: $k_{\mathrm{on}} = k_{\mathrm{on}}^{0} \frac{l_{\mathrm{bind}}}{Z} e^{\left(-\frac{k(\delta - l_{\mathrm{b}})^2}{2 k_{\mathrm{B}} T}\right)}$, where $k_{\mathrm{on}}^{0}$ is the intrinsic association rate. The partition function can be written as $Z = \sqrt{\frac{\pi k_{\mathrm{B}} T}{2k}} \left[ \mathrm{erf}\left( (\delta - l_{\mathrm{b}}) \sqrt{\frac{k}{2 k_{\mathrm{B}} T}} \right) + \mathrm{erf}\left( l_b \sqrt{\frac{k}{2 k_B T}} \right) \right]$, erf representing the error function.

**4. The Elastic-Stochastic model**

To capture the stochastic bond formation and disruption during virus invasion, the kinetic Monte Carlo (KMC) method was used with the defined biochemical reaction rates [4]. The core assumption in KMC is that the probability of any event occurring is proportional to its reaction rate. In the virus invasion, spike–ACE binding and dissociation, and S1/S2 subunit disengagement are treated as independent biochemical reaction events. The values of the corresponding reaction rates, $k_{\mathrm{on}}$, $k_{\mathrm{off}}$, and $k_{\mathrm{dis}}$ are determined based on the force distribution among the bound bonds and the surface separation at spike–ACE2 protein pairs, obtained by the elasticity analysis. By summing up the chemical reaction rates of all possible events in the system, a total reaction rate, $k_{\mathrm{tot}}$, is obtained. The KMC algorithm is then performed to simulate the stochastic dissociation and association of bonds during the invasion. At each moment, a random number, $\delta_1$, between 0 and 1, is generated to decide what event is occurring. Another random number, $\delta_2$, between 0 and 1, is generated to determine the duration of this event by the following equation: $\mathrm{d}\tau = -ln(\delta_2)/k_{\mathrm{tot}}$. By integrating the KMC method into our model, we can effectively simulate the intricate and stochastic nature of the bond formation during virus invasion. Following the event, the system proceeds to an elastic update: the shape of the cell membrane and the forces applied to each spike–receptor bond are recalculated based on the new binding configuration. This updated mechanical state feeds back into the next KMC step, maintaining full coupling between stochastic bond kinetics and mechanical deformation. All simulations begin with a viral particle resting on a flat, undeformed cell membrane, with no spike–receptor bonds initially present. By iteratively solving this coupled elastic–stochastic problem, the model captures the dynamic interplay between bond formation, dissociation, membrane bending, and force regulation throughout the viral entry process. This framework allows for single-bond resolution tracking of the invasion pathway under various mechanochemical conditions.

A representative set of parameters corresponding to the WT SARS-CoV-2 variant was used in our simulations (Table S1), and this serves as the baseline for evaluating variant-specific effects on invasion dynamics.

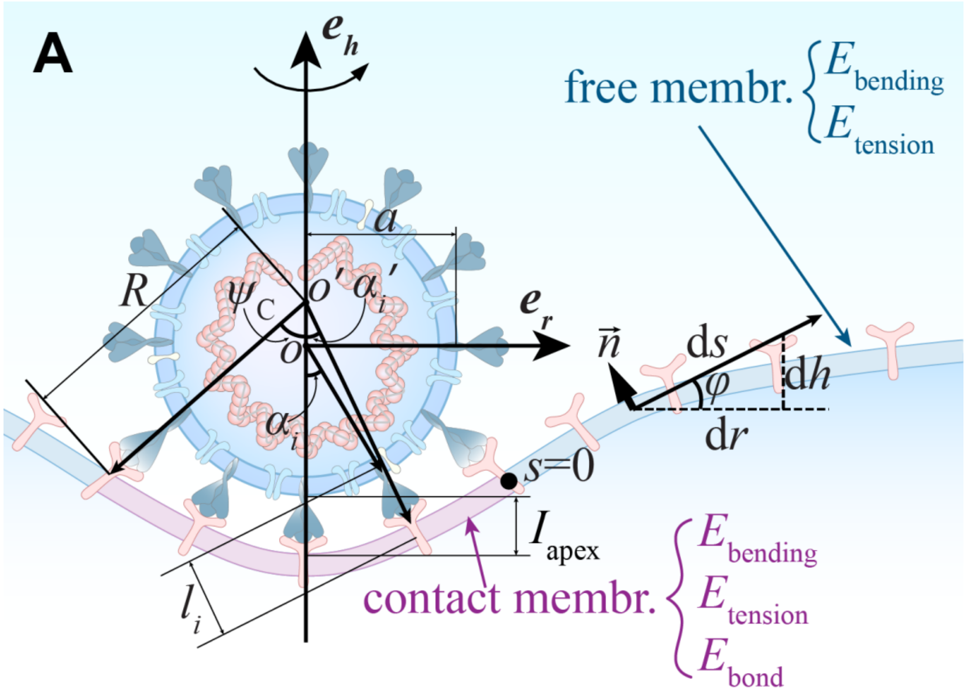


**Figure S1.** The coordination definition and the geometric parameters in the analytic mechanical model.

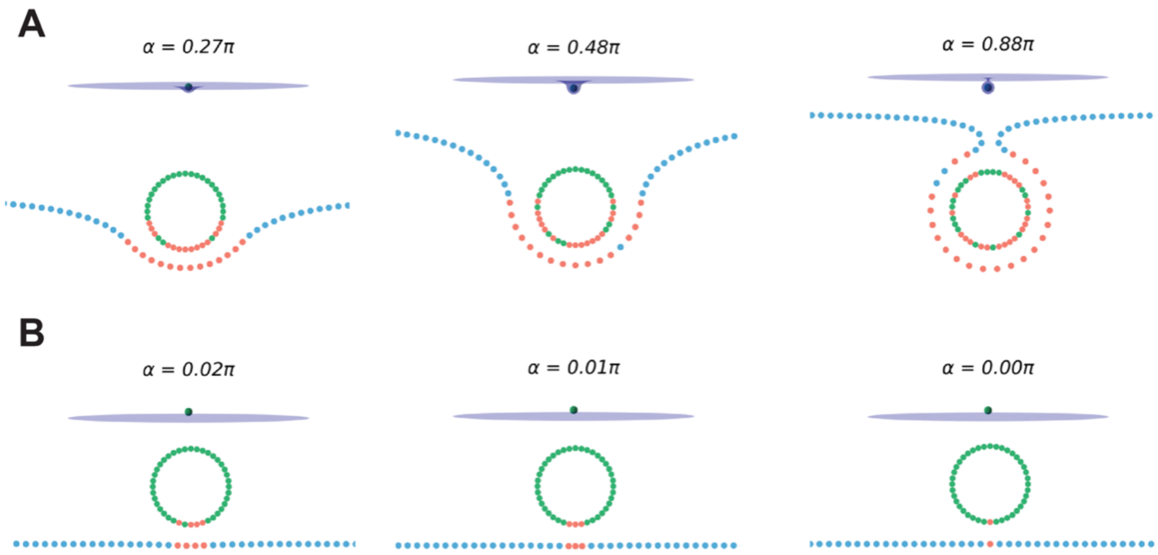


**Figure S2.** The snapshots of the virus invasion at varying wrapping angles for two distinct bond behaviors: a catch bond scenario with $k_C/k_S = 100$ (**A**) and a slip bond scenario with $k_C/k_S = 1$ (**B**). The top and bottom panels display a 3-D and 2-D view of the virus, respectively. In the 2-D view, the spike and ACE proteins that have not formed bonds are depicted in green and blue, respectively, while those that have formed bonds are highlighted in red.

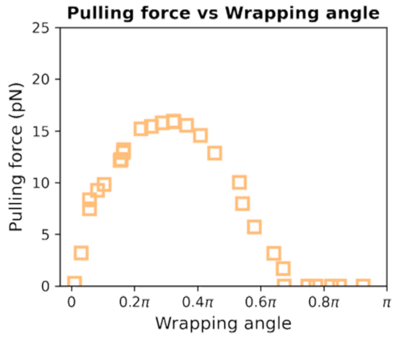


**Figure S3.** The representative pulling force at the wrapping edge during virus invasion.

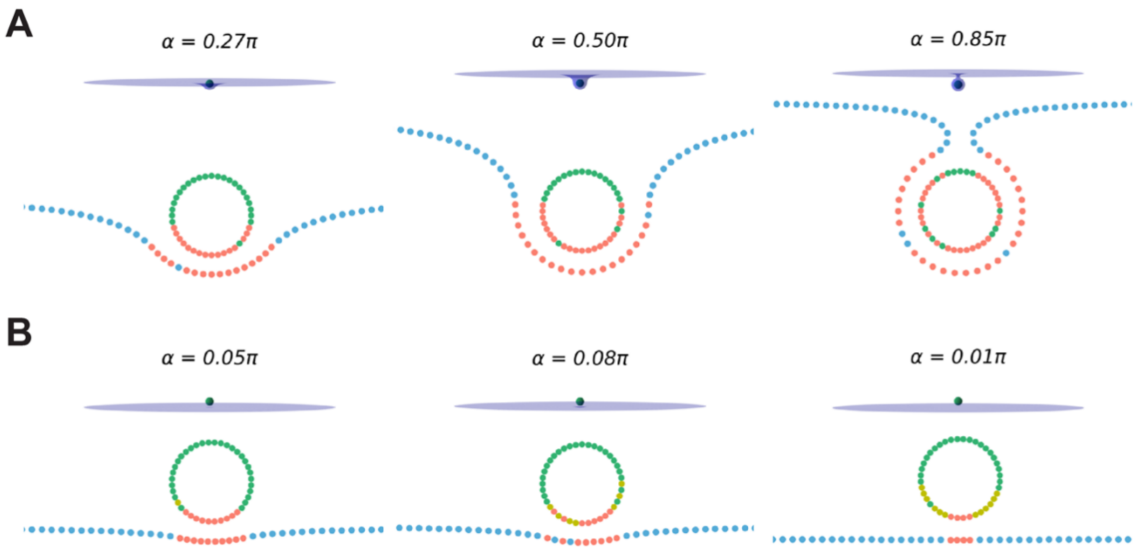


**Figure S4.** The snapshots of the virus invasion at varying wrapping angles for two distinct force acceleration on S1/S2 disengagement: a slow acceleration scenario with d$x$ = 0.5 nm (**A**) and a fast acceleration scenario with d$x$ = 4 nm (**B**). The top and bottom panels display a 3-D and 2-D view of the virus, respectively. In the 2-D view, the blue, green, and red dots share the same definition as in **Figure S2**, and the yellow dots represent the spikes, in which the S1/S2 has successfully disengaged.

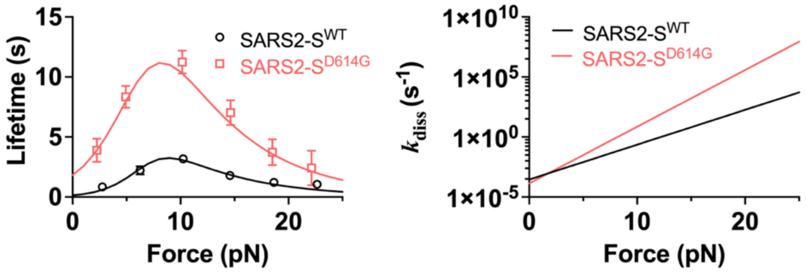


**Figure S5.** Fitting to the force-regulated spike–ACE2 dissociation (**A**) and S1/S2 disengagement (**B**) of WT and D614G variants.

**Table S1.** Summary of parameters used in the simulations.

| | Parameter | Value | Unit | Reference |
|---|---|---|---|---|
| Bending Rigidity | $\kappa$ | 80 | pN·nm | 1,7,14 |
| Stiffness Parameter | $k$ | 2 | pN/nm | 1 |
| Radius of Virus | $a$ | 40 | nm | 1 |
| Number of Spikes On Virus Surface | $N$ | 40 | | 1 |
| Density of Ace2 Receptor on Membrane | $\xi_r$ | 0.01 | $nm^{-2}$ | 1 |
| | $k_B T$ | 4.2 | pN·nm | |
| Resting Length | $l_b$ | 23 | nm | 1 |
| Surface Tension | $\sigma$ | 0.01 | pN/nm | 7,14 |
| Binding Rate | $k_{on}^{0}$ | 2 | $s^{-1}$ | 5,15 |

**Table S2.** VOCs Bell model parameters for S1/S2 disengagement

| | WT | D614G |
|---|---|---|
| $k_0$ ($s^{-1}$) | 0.000287 | 0.000136 |
| d$x$ (nm) | 2.7667 | 4.5035 |

**Table S3.** VOCs one-state two-pathway catch bond model parameters for spike-ACE2 dissociation

| | WT | D614G |
|---|---|---|
| $x_C$ (nm) | -2.16 | -1.54 |
| $x_S$ (nm) | 0.58 | 0.59 |
| $k_C$ ($s^{-1}$) | 6.66 | 0.53 |
| $k_S$ ($s^{-1}$) | 0.07 | 0.02 |

**Video S1.** The movie of the virus invasion for two distinct bond behaviors: a catch bond scenario with $k_C/k_S$ = 100 (upper) and a slip bond scenario with $k_C/k_S$ = 1 (lower).

**Video S2.** The movie of the virus invasion for two distinct force acceleration on S1/S2 disengagement: a slow acceleration scenario with d$x$ = 0.5 nm (upper) and a fast acceleration scenario with d$x$ = 4 nm (lower).

#### Supplementary Material References
